# Evaluating evidence for co-geography in the *Anopheles-Plasmodium* host-parasite system

**DOI:** 10.1101/2023.07.17.549405

**Authors:** Clara T. Rehmann, Peter L. Ralph, Andrew D. Kern

## Abstract

The often tight association between parasites and their hosts means that under certain scenarios, the evolutionary histories of the two species can become closely coupled both through time and across space. Using spatial genetic inference, we identify a potential signal of common dispersal patterns in the *Anopheles gambiae* and *Plasmodium falciparum* host-parasite system as seen through a between-species correlation of the differences between geographic sampling location and geographic location predicted from the genome. This correlation may be due to coupled dispersal dynamics between host and parasite, but may also reflect statistical artifacts due to uneven spatial distribution of sampling locations. Using continuous-space population genetics simulations, we investigate the degree to which uneven distribution of sampling locations leads to bias in prediction of spatial location from genetic data and implement methods to counter this effect. We demonstrate that while algorithmic bias presents a problem in inference from spatio-genetic data, the correlation structure between *A. gambiae* and *P. falciparum* predictions cannot be attributed to spatial bias alone, and is thus likely a genetic signal of co-dispersal in a host-parasite system.

## Introduction

A nearly ubiquitous observation in biology is that individuals within a species that live closer together in space are more closely related genetically than individuals that live further apart. Such isolation by distance, as it was referred to by Wright (1943), and its fundamental determinant – the spatially limited dispersal of individuals across a landscape – can have profound impacts on patterns of genetic variation within and between populations (Whitlock and McCauley, 1999; Bradburd and Ralph, 2019), genetic association studies (Mathieson and McVean, 2012; Battey et al., 2020b), and the potential for local adaptation (Barton, 2001). As befits the central importance of understanding dispersal, over many decades much work has gone in to trying to estimate migration of individuals between local demes or across continuous space (Rousset, 1997; Marcus et al., 2021).

Combining spatial and genetic data can reveal a wealth of information about biological and ecological processes: for example, these data can be used to predict a genetic sample’s location (Wasser et al., 2004; Finch et al., 2020; Guillot et al., 2015), estimate dispersal and migration patterns (Marcus et al., 2021; Smith et al., 2023), or describe major features of spatial genetic structure (Bradburd et al., 2013). Indeed with the increasing availability of geo-referenced genotypic data as well as increasingly sophisticated methods of spatial population genetics simulation (Battey et al., 2020b), there is growing opportunity to explore the potential of identifying biological processes through patterns of isolation by distance. Recently, Battey et al. (2020a) introduced Locator, a state-of-the-art, deep learning method for predicting the geographic origin of a genetic sample. Locator is able to achieve extremely accurate prediction on large, publicly available datasets. Further, Battey et al. (2020a)’s analysis indicated that errors in geographic prediction of locations (which we refer to as *residuals*) can be informative of those individuals’ geographic ancestry: for example, when applied to human data, predicted locations of South African Bantu individuals trended towards west Africa, the historic origin of the Bantu Expansion.

More generally, we would expect various biological processes to affect Locator’s residuals, and so may be able to examine residuals to infer the processes that give rise to these patterns. For instance, two processes we expect to affect residuals are heterogeneous dispersal and range expansion. If dispersal is heterogeneous across the landscape, with high dispersal in some regions and low dispersal in others, regions with higher levels of gene flow would have less accurate geographic prediction. In the case of range expansions, one would expect the expanding lineage to genetically resemble individuals from the original range - a prediction we see reflected in the South African Bantu example. In both cases, though we may be unable to identify individual instances of dispersal, errors in Locator’s predictions can be interpreted to provide clues about underlying population processes.

Battey et al. (2020a) applied Locator to data from the malaria vector *Anopheles gambiae* (Ag1000G; Anopheles gambiae 1000 Genomes Consortium (2017)) and its associated parasite *Plasmodium falciparum* (Pf7k, Pearson et al. (2019)), and were able to predict geographic origin of individuals to within a median error of 16.9 km and 85 km, respectively. We were interested in exploring if any structure might be present in the residuals from Locator’s predictions. In particular, close examination of patterns in prediction error between species on the African continent suggested to us that perhaps residuals between spatially proximate individuals from the *Anopheles gambiae* and *Plasmodium falciparum* datasets might be correlated. Given that we expect Locator’s residuals to be affected by heterogeneous gene flow, these errors may reflect shared non-uniform migration patterns between host and parasite, for instance a broad corridor of dispersal throughout West Africa but more restricted gene flow in Central and Eastern parts of the continent.

Seen from the standpoint of host-parasite ecology, such correlations in the residuals of geographic predictions may not be surprising. If parasites depend on hosts for dispersal, then we might indeed expect a close correspondence between the spatial genetic structure of these interacting species. Such patterns have been observed before: for instance, Small et al. (2019) found a tight correspondence between divergence times between geographical locations of the filarial nematode parasite *Wuchereria bancrofti* and known instances of human migration, thus linking parasite divergence to (human) host dispersal. At the limit of low parasite dispersal and/or considered over longer evolutionary time scales, these dynamics can translate into co-speciation, where patterns of splitting in host and parasite closely mirror one another (e.g., Moran and Baumann, 1994; Clayton et al., 2003; Jousselin et al., 2008), and many phylogenetic-based methods have been developed to identify patterns of host-parasite cospeciation (e.g., Hafner and Nadler, 1988; Huelsenbeck et al., 1997, 2000) on a macroevolutionary scale.

While residuals from Locator’s predictions may reflect geographic patterns of gene flow in this case common dispersal patterns between *A. gambiae* and *P. falciparum* populations the data used to train the network does not represent the complete geographic and genetic diversity of these populations. Indeed in many biological datasets, the geographic locations of samples do not capture an entire landscape and instead are often biased, for instance towards easily sampled locations; this inconsistency in spatial sampling is a well-documented problem in applications such as ecological niche modeling (Kramer-Schadt et al., 2013). Further, it is now well appreciated in the broader machine learning field that models trained from even subtly biased training sets, will yield predictions that reflect those biases with increasingly broad societal implications (e.g., Buolamwini and Gebru, 2018; Mehrabi et al., 2021). There is no reason to believe that such biases would not be present in our own spatial population genetic predictions.

In this study we explore the evidence of correlation between Locator’s geographic predictions of *A. gambiae* and *P. falciparum* in sub-Saharan Africa, and investigate whether these patterns emerge from spatial imbalance in training data. Using extensive continuous space forward-in-time population genetic simulations, we demonstrate that non-uniform spatial sampling in training data is a pernicious problem in geographic prediction, but does not alone account for the residuals observed in *A. gambiae* and *P. falciparum* populations. We explore and implement methods to counter the bias created by non-uniform training sets, and demonstrate conditions under which we might expect geographic prediction to be more or less biased. Applying these methods to our predictions of *A. gambiae* and *P. falciparum* locations, we show that while non-uniform sampling in our training sets may elevate correlations between residuals, our signal is beyond what would be expected from sampling considerations alone. We hypothesize that the correlated patterns between host and parasite residuals demonstrate the evolutionary consequences of linked dispersal between *A. gambiae* and *P. falciparum* on a short timescale.

## Results

### Errors in geographic predictions of *A. gambiae* and *P. falciparum* are correlated

*A. gambiae* mosquitoes are the vector of the malaria-causing parasite *P. falciparum*. While the parasite spends most of its life cycle in humans, female mosquitoes acquire *P. falciparum* gametocytes when feeding on an infected human, which then sexually reproduce before being transmitted as fertilized sporozooites to a human host during the next blood meal (Hall et al., 2005). Though per generation dispersal is limited to perhaps as little as roughly six kilometers per generation (Smith et al., 2023), the dispersal of infected *A. gambiae* mosquitoes will consequently bring along the infecting *P. falciparum*. It is less clear whether this host’s dispersal meaningfully affects *P. falciparum* movement and genetic structure.

When applying Locator to empirical data from *A. gambiae* and *P. falciparum*, we observed similar patterns in prediction errors (i.e. residuals) between spatially proximate samples of the host and its parasite (Figure 1 A, B, C). Battey et al. (2020a) showed that Locator predictions reliably reflect individuals’ geographic ancestries in some well-documented cases from humans. Thus we hypothesize that our observation of similar prediction errors between between host (*A. gambiae*) and parasite (*P. falciparum*) may reflect common, recent co-geographic patterns. These patterns are driven primarily by large prediction errors in West Africa, potentially representing a region of relatively high gene flow for both host and parasite. Essentially because *P. falciparum* depends on *A. gambiae* mosquitoes for a portion of its life cycle, some degree of dispersal will be due to dispersal of infected mosquitoes, which should induce co-variation in their evolutionary histories across the landscape.

**Figure 1:**
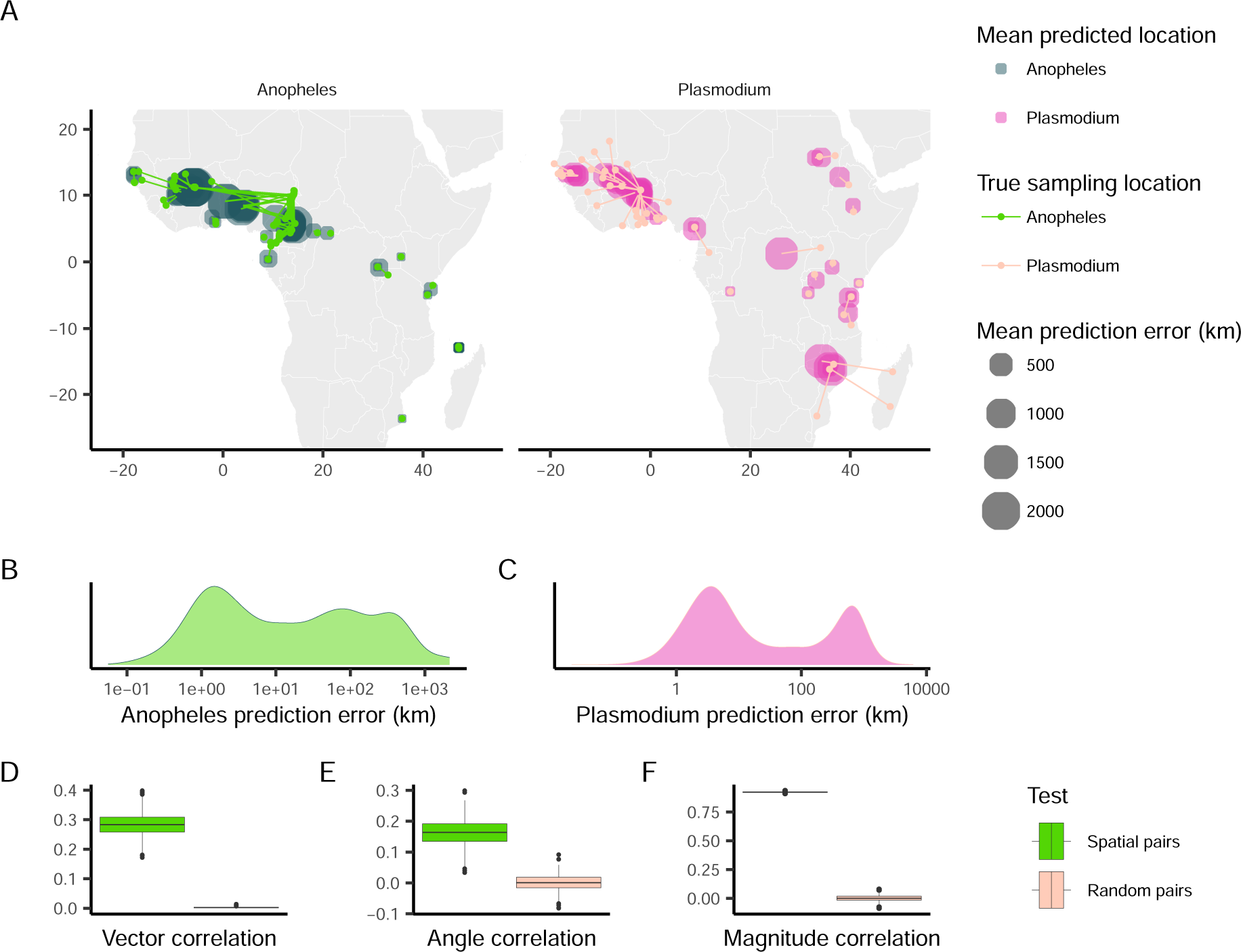
Predicted locations of *Anopheles gambiae* and *Plasmodium falciparum* samples. (A) Geographic centroids of predictions for each unique sampling site. True sampling locations are connected to their mean predicted location; the size of predicted locations represents error in km. (B, C) Distribution of error magnitudes for *A. gambiae* and *P. falciparum* predictions. (D, E, F) Vector, angle, and magnitude correlations between spatially paired and randomly paired *A. gambiae* and *P. falciparum* residuals.

We first aimed to quantify the relationship between the two datasets’ residuals by applying Crosby et al. (1993)’s technique for measuring correlation between pairs of vector-valued time series. Originally developed for geophysical applications, this vector correlation metric accounts for both magnitude and direction of a set of vectors, and ranges from zero (no correlation) to two (complete correlation). To apply this method to our datasets, we tested for correlation between paired *A. gambiae* and *P. falciparum* individuals from the same approximate geographic area, defining vectors from an individual’s true location to their predicted location (i.e. the residuals of our predictions). Because multiple samples can occur in one location, we ran this analysis multiple times, with each iteration resulting in different pairings of spatially proximate individuals. For comparison, we also permuted spatial pairings between species to create random pairs with respect to space, and then applied the same correlation procedure to those randomized datasets. This set of randomly paired data represents a naive null distribution to which we compare our observed correlations.

The mean vector correlation statistic between spatially paired residuals was 0.284 (*SD* = 0.038), which was significantly different (Wilcoxon test, *p <* 2.2 *×* 10*^−^*^16^) from the correlation observed in our permuted pairs (Figure 1 D). This suggests that there are common patterns of error in Locator’s predictions of *A. gambiae* and *P. falciparum* locations. However, Crosby et al. (1993)’s vector correlation statistic can be influenced by both the angles and magnitudes of the vectors in question, making interpretation hazy. Because of this, we also performed separate Spearman rank correlation tests on the angle and magnitude of residual vectors individually from the same spatially- and randomly-paired datasets (Figure 1 E, F). Both the angles and magnitudes of error vectors were significantly correlated between spatially-proximate *A. gambiae* and *P. falciparum* samples (*R* = 0.163 (*SD* = 0.041)*, p <*2.2 *×* 10*^−^*^16^ and *R* = 0.924 (*SD* = 0.004)*, p <*2.2 *×* 10*^−^*^16^), further supporting that geographic errors in predictions are similar between the two species.

We further examined the correlation in genetic structure between the species by comparing principle components of species’ genotype matrices (Figure S3). This reveals a similar pattern whereby spatially proximate samples are embedded in PCA space in a correlated fashion. Indeed, longitudinal differention seems to be shared among *A. gambiae* and *P. falciparum*.

### Is this a biological signal?

The observed correlation between *A. gambiae* and *P. falciparum* residuals is beyond what we would expect under a naive null, particularly in West Africa, but does this implicate the movement of mosquito vectors in the spatial spread of malaria parasites? Or a genetic signal of shared geographic history between these species? Further, while our observed residuals align with known corridors of gene flow, particularly in West Africa (Anopheles gambiae 1000 Genomes Consortium, 2017), they also trend towards locations with increased sampling density. This raises a possible statistical issue – are our predictions simply pointing towards geographic regions that are over-represented in our training set? Unbalanced training data is well-known to create issues for machine learning based inference, so we were interested to assess how geographic prediction is influenced by training data that is non-uniformly sampled throughout space and if (or to what degree) such an effect might be responsible for our observed correlation between *A. gambiae* and*P. falciparum* residuals.

### Spatial bias in training data leads to biased geographic predictions

To understand the impact of spatially imbalanced training data on predictions, we applied Locator to samples from populations simulated using a continuous space, forward-in-time model implemented in SLiM (Haller and Messer, 2019). In these simulations, the spatial aspect of dispersal, competition, and mating of individuals are each controlled by a Gaussian kernel with mean zero and standard deviation *σ* in each dimension, which we vary across simulations.

To generate spatially imbalanced data from these simulations, we created training sets where samples were increasingly likely to be sampled from the right half of the landscape (Figure 2 A). In these simulations, the relatedness between individuals varies smoothly across the landscape (i.e. the degree of spatial population structure is relatively continuous (Battey et al., 2020b)), so we expect any systematic spatial patterns in prediction error to be attributed to sampling imbalance in training data. After training Locator on 450 individuals sampled in this fashion, we then used the trained network to predict the locations of held-out individuals that were sampled uniformly across the landscape. By separating the vector formed between an individual’s true and predicted location into its *x* and *y* components, we were able to separately quantify error along the axes of imbalanced sampling (*x*) and random sampling (*y*). These values were then averaged across each Locator run, summarizing the trends in prediction errors for a given training set.

**Figure 2:**
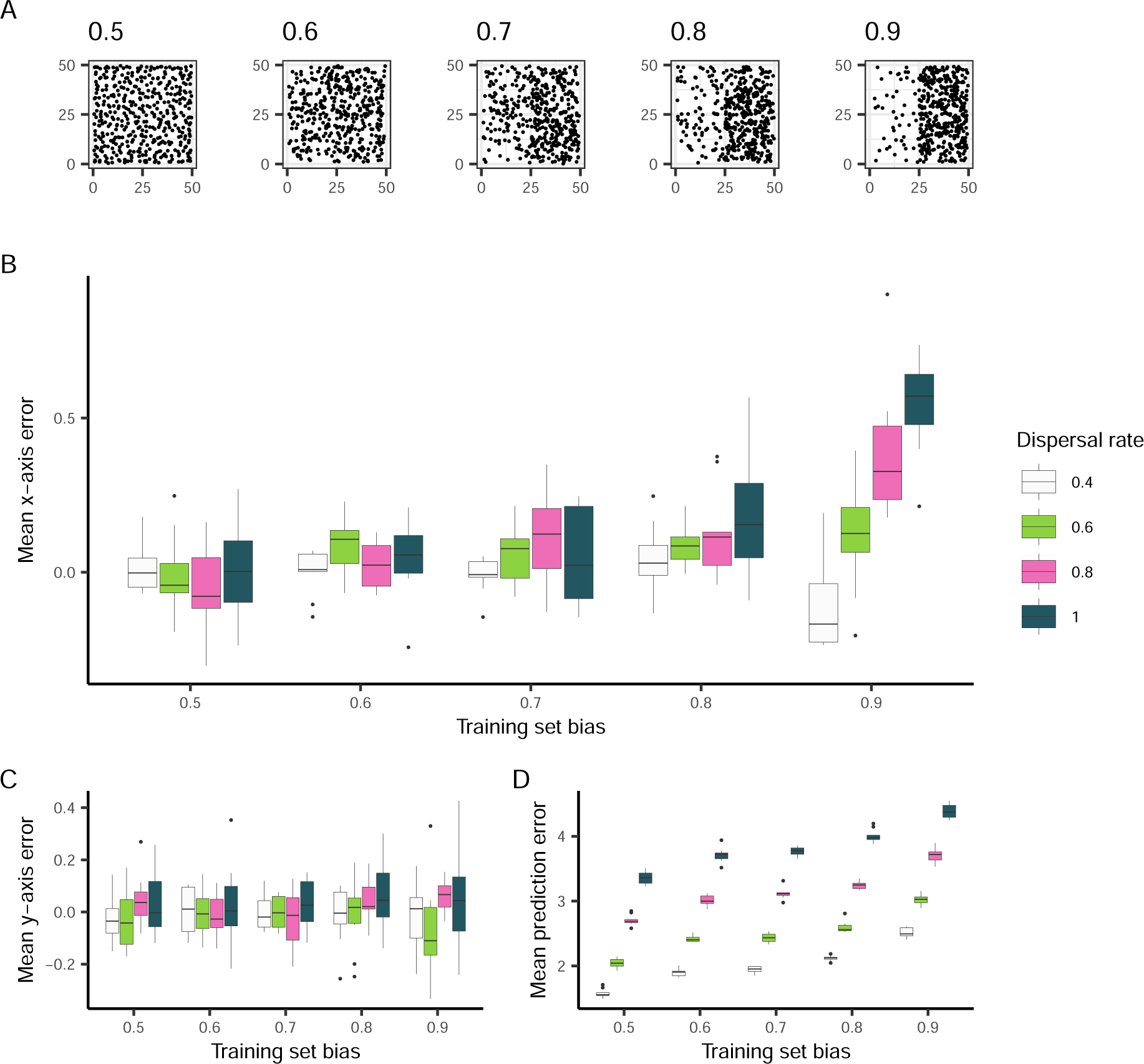
The effect of training set bias on Locator residuals. (A) Training sets sampled with increased bias towards the right side of the landscape. (B) Mean per-run prediction error along the x axis as a function of training set bias, (C) Mean per-run prediction error along the y axis, (D) Mean per-run error magnitude.

We found that when Locator is trained with spatially biased data, predictions are biased towards more densely sampled regions of the landscape. In other words, training sets that were biased towards the right half of the landscape resulted in an excess of predictions to the right of samples’ true locations, with this effect becoming more pronounced as dispersal rate (*σ*) increases (Figure 2 B). We quantified the contribution of training set bias and per-generation dispersal rate (both normalized between 0 and 1) to x-axis error using multiple linear regression; both variables have a significant positive effect on x-axis error (*β*_1_ = 0.128*, p* = 1.61 *×* 10*^−^*^15^ and *β*_2_ = 0.090*, p* = 1.17 *×* 10*^−^*^8^ for bias and *σ*, respectively).

While bias in prediction errors is observed along the axis of imbalanced sampling, there is no relationship between y-axis error and training set bias (Figure 2 C), suggesting that bias in residuals reflects the spatial imbalance in training data. A linear model confirms that training set bias has no effect on y-axis error, (*p* = 0.244) but dispersal rate has a small but significant positive effect (*β*_2_ = 0.039*, p* = 0.013). This phenomenon is likely due to overfitting: when learning the relationship between genotype and geography, Locator aims to minimize the distance between true and predicted locations of training data, which, with biased training sets, may be most efficiently done by learning to predict towards densely-sampled regions of the landscape, regardless of true sample location. This prediction bias is amplified at higher dispersal rates due to the reduction in spatial genetic structure that Locator is picking up on, and predictions are increasingly based on locations the network has already seen.

### Spatial distribution of training data does not explain Anopheles residuals

To assess the contribution of spatial sample imbalance to observed patterns in empirical residuals, we again applied Locator to simulated randomly dispersing populations, this time using training samples that mimic the spatial distribution of the Ag1000G dataset. To accomplish this we performed simulations which reflect Africa’s *Anopheles gambiae* population, which has an estimated per-generation dispersal rate of 13 kilometers per generation and an effective population size of approximately 1 million individuals (Smith et al., 2023; Anopheles gambiae 1000 Genomes Consortium, 2017). Populations were simulated across a landscape representing Sub-Saharan Africa, which contains all sampling sites and reflects the general range of *Anopheles* mosquitoes. After running these simulations, we sampled individuals in the same spatial pattern as the Ag1000G dataset and applied Locator in a windowed analysis across a 10^8^ bp genome, using 2Mbp windows that capture roughly 80, 000 SNPs each - less genetic information than the empirical data. A rotating 10 percent of samples were held out during training runs and used as a prediction set in the same fashion as our empirical analysis. The outcome of these simulations is a rough approximation of African *A. gambiae* populations under neutral spatial dynamics, not accounting for variation in population density, seasonal changes, or mosquito-host dynamics (among other factors). However, we can use these simulations to represent a baseline estimate of population structure across space under random dispersal. As in our right-biased training sets from randomly dispersing simulations, we can then assess the impact of spatially imbalanced sampling on Locator’s predictions, this time comparing them to predictions from the true Ag1000G dataset and the data from *P. falciparum*.

Despite training data being sampled unevenly from the landscape, Locator’s performance on these simulated populations was relatively accurate (Figure 3). While predictions for individuals near landscape edges were slightly biased towards the center, predictions near the center of the landscape - particularly in West Africa - were surprisingly close to individuals’ true locations. This is in contrast to Locator’s performance on the empirical *A. gambiae* dataset, where samples near the edges of the continent are predicted accurately and West African samples have large, inward-pointing residuals (Figure 3A). This increase in prediction accuracy is also visible in the distribution of error magnitudes, which is bimodal in the case of empirical data but more evenly distributed in the case of the simulated data (Figure 3B). We then tested for correlation between empirical *P. falciparum* and simulated *A. gambiae* residuals. The goal of this comparison in essence is to remove any possibility of shared or biased dispersal histories between species while still allowing for continuous population structure. We find that vector and angle correlations when testing against simulated data are significantly less that what we observe in the real data, though a correlation remains in the magnitude of residuals (Figure 3C, D, E).

**Figure 3:**
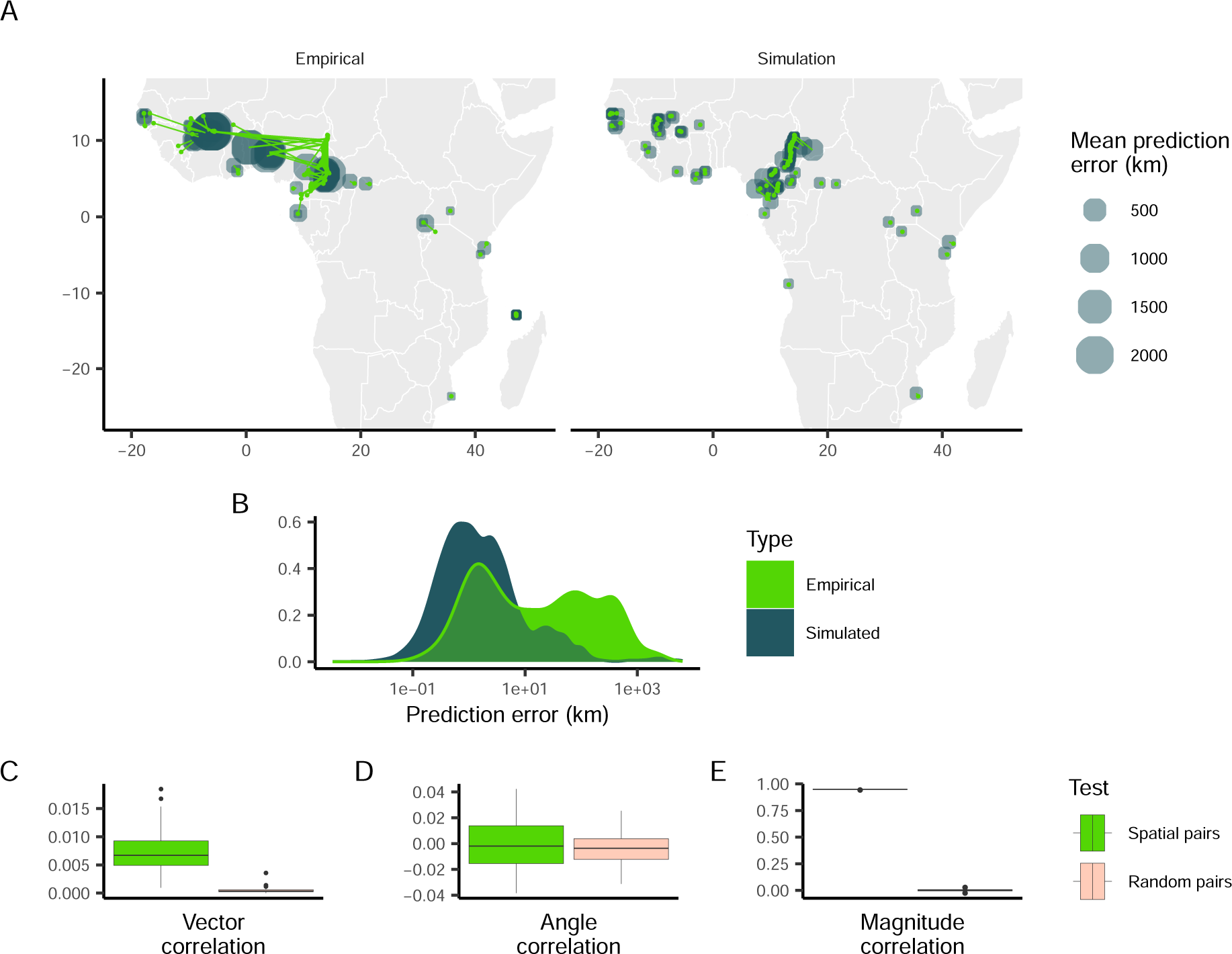
Predicted locations of empirical and simulated *Anopheles gambiae* samples. (A) Geographic centroids of predictions for each unique sampling site. True sampling locations are connected to their mean predicted location; the size of predicted locations represents error in km. (B) Distribution of error magnitudes. (C, D, E) correlation coefficients between simulated Anopheles and empirical Plasmodium predictions for the residual vectors, residual angles, and residual magnitude respectively.

While the correlation between empirical *P. falciparum* and simulated *A. gambiae* error magnitude indicates that the spatial distribution of samples may influence similarities in predictions, many major features of the empirical predictions of *A. gambiae* locations are not reproduced by simulated neutral spatial processes. In particular, residuals of empirical samples in West Africa are large and trend towards the center of the region in a similar fashion to *P. falciparum*, while residuals of simulated samples are relatively small. *A. gambiae* populations in this region have high levels of gene flow, despite large geographic distances, while populations in Central Africa show more genetic differentiation (Anopheles gambiae 1000 Genomes Consortium, 2017; Clarkson et al., 2020). These results suggest that the correlation between prediction errors observed when Locator is applied to the Ag1000G and Pf7k datasets are not only due to the spatial distribution of sampling locations, and may be indicative of common gene flow and non-uniform migration patterns.

### Accounting for spatially biased data in geographic prediction

While our simulations approximating *A. gambiae* populations suggested that patterns in Locator’s prediction errors were not wholly due to spatial bias in empirical datasets, we were interested in reducing Locator’s prediction bias when applied to spatially biased datasets. Because most natural populations are not sampled in a way that accurately reflects their range and density, it is possible that spatial sampling bias may impact Locator’s predictions in other empirical applications. To see if prediction bias could be reduced in these situations, we again applied Locator to our simulation data. Because the relationship between training set and prediction bias is stronger at higher dispersal rates, we focused on simulations with a dispersal rate of *σ* = 1.0 per generation. We first attempted to reduce prediction bias by downsampling individuals in densely-sampled regions of the landscape - this successfully reduced mean prediction bias (though not significantly), but in turn increased prediction error (Figure 4 A, C, D, E). This makes sense, as downsampling reduces the overall amount of training data available. In an attempt to make use of all the samples in our dataset while still accounting for spatial bias, we then implemented a sample weighting scheme in Locator’s training process.

**Figure 4:**
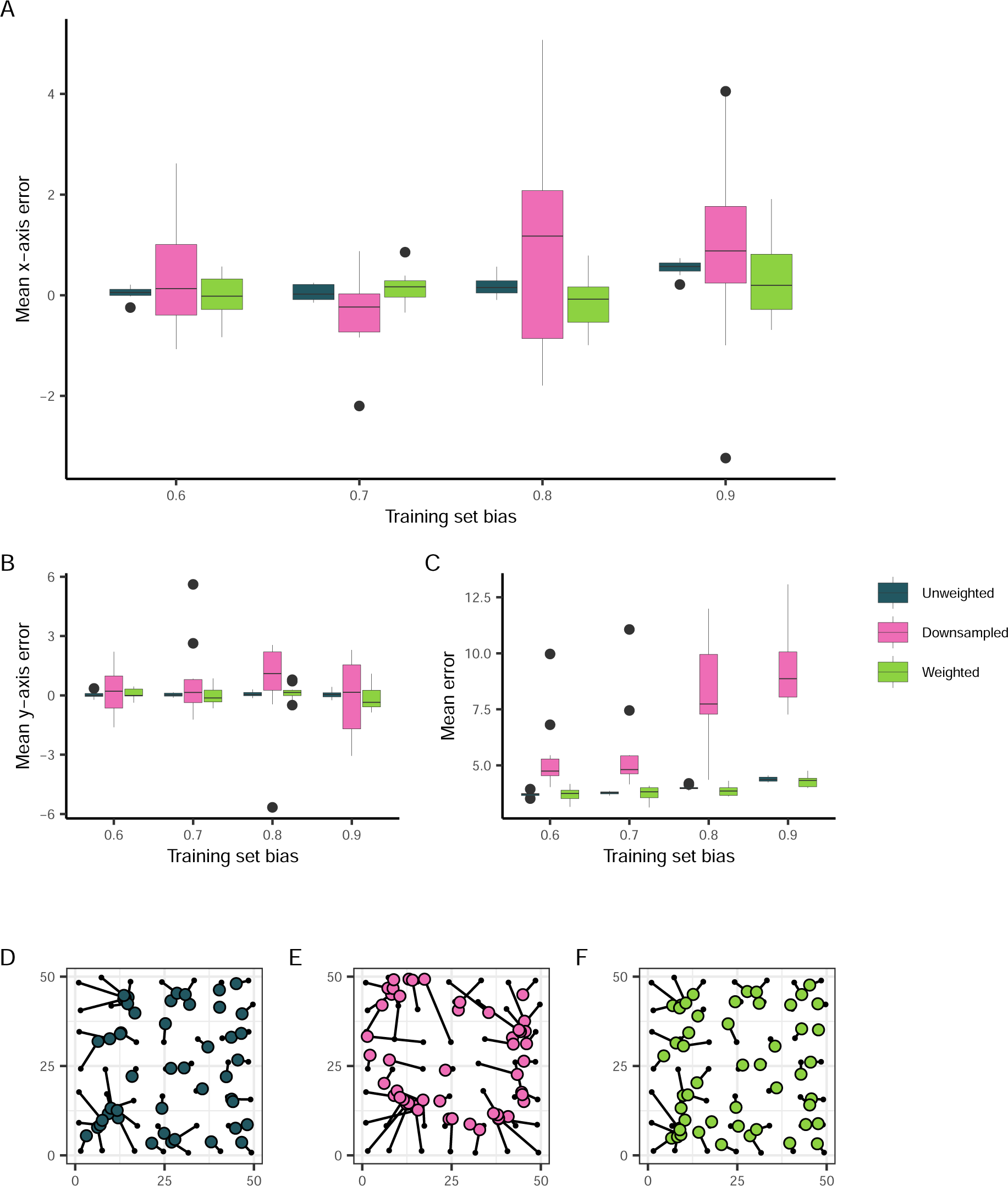
The effect of training set bias using sampling and training modifications. (A) Mean per-run prediction error along the x axis as a function of training set bias, (B) Mean per-run prediction error along the y axis, (C) Mean per-run error magnitude. (D, E, F) Example predictions at bias = 0.9 for unmodified, downsampled, and weighted training sets, respectively. True sample locations (black points) are connected to their predicted location (colored points).

During each epoch of training, Locator aims to minimize the sum of distances between true and predicted locations (the *loss*). In the case of spatially biased datasets, this is done most efficiently by learning to predict in areas with dense sampling, leading to prediction bias. To reduce bias towards densely sampled portions of the landscape during training, we weighted each sample’s contribution to the overall loss according to its local spatial density: samples in sparse regions were upweighted, and samples in dense regions were downweighted. To calculate sample weights we fit a kernel density estimate over the landscape, then assigned each sample a weight value inversely proportional to the sampling density in that region raised to a power *λ*. We varied two parameters when assigning sample weights: bandwidth, which controls the “smoothness” of the kernel density estimate (Figure S6 B), and *λ*, which effectively increases the difference between the lowest and highest weights (Figure S6 A). Further discussion of sample weight parameterization is given in the Methods (section “Sample Weighting”). As kernel bandwidth and *λ* are effectively hyper-parameters of our model used to tune weighting, we performed a grid search across a discretized space of *λ* and bandwidth to identify effective parameter combinations that reduced loss for a given training set.

In application, it is difficult to disentangle biological causes of prediction error from potential bias caused by a sampling scheme. Therefore in our grid search we aimed to identify the parameter combination that minimized *overall* prediction error as opposed to prediction bias (Figure S5). The parameter combination that most significantly reduced error on training sets with a sampling bias of 0.9 was a bandwidth value of 0.25 and a *λ* value of 10*^−^*^5^ which reduced median prediction error by 14% (from 4.374 to 3.753 units). Examining prediction bias along the x axis, this reduces the linear relationship between normalized bias and x-axis error from 0.3838 (*p <* 2 *×* 10*^−^*^16^) to 0.1473 (*p* = 0.1449) (Figure 4 A). Like downsampling, sample weighting also results in a bias-variance tradeoff, though less dramatically: for a training set bias of 0.9, sample weighting reduced a run’s mean x-axis error from 0.5398 to 0.2947 units, but increased the standard deviation in x-axis error from 0.1529 to 0.8224 units (Figure 4 A, C).

Overall, weighting sample importance during training provides a way to reduce, though not completely eliminate, the issues caused by spatially biased training datasets. Methods to calculate and implement sample weights during training are now included in Locator, and future applications of the neural network to empirical datasets may benefit from considering the spatial distribution of available training samples during analysis.

### Correlation between *A. gambiae* and *P. falciparum* prediction errors is not due to bias

Having identified a method of reducing prediction bias when training with spatially uneven data, we then used sample weights when applying Locator to the Ag1000G and Pf7k datasets. To identify parameter values for assigning sample weights, we performed a grid search over bandwidth values between 1 and 10,000 and *λ* values between 1 and 10. To reduce compute time, we ran Locator on 100,000 randomly chosen SNPs from chromosome 2R for the *A. gambiae* data and on the first 500kb of chromosome 0 for the *P. falciparum* data, then used the final validation loss after training to assess performance. For both datasets, a combination of bandwidth = 100 and *λ* = 1.0 resulted in the largest decrease in validation loss. The mean validation loss for the unweighted *A. gambiae* dataset was 0.115 *±* 0.018; with sample weighting, the mean validation loss was reduced to 0.098 *±* 0.012. For the unweighted *P. falciparum* dataset, the mean validation loss was 0.351 *±* 0.039; with sample weighting, this was reduced to 0.297 *±* 0.026.

We then ran Locator on the full *A. gambiae* and *P. falciparum* datasets using sample weights, setting bandwidth = 100 and *λ* = 1.0 (Figure 5 A). This only reduced mean prediction error in the *A. gambiae* dataset from 130.3 to 119.8 km (not significant by Wilcoxon test), and slightly elevated median prediction error from 12.8 to 13.4 km. In the *P. falciparum* dataset, mean prediction error was significantly reduced from 203.1 to 151.2 km (Wilcoxon test, *p* =2.24 *×* 10*^−^*^14^), and median prediction error was reduced from 8.2 to 6.9 km (Figure 5 B, C). With sample weighting, the correlation between spatially proximate *A. gambiae* and *P. falciparum* residuals is still significant, though reduced in magnitude (Figure 5 D, E, F): correlation between vectors is *R* = 0.079 *±* 0.033, correlation between residual angles is *R* = 0.076 *±* 0.037, and correlation between error magnitudes is *R* = 0.907 *±* 0.006 (all metrics *p <*2.2 *×* 10*^−^*^16^).

**Figure 5:**
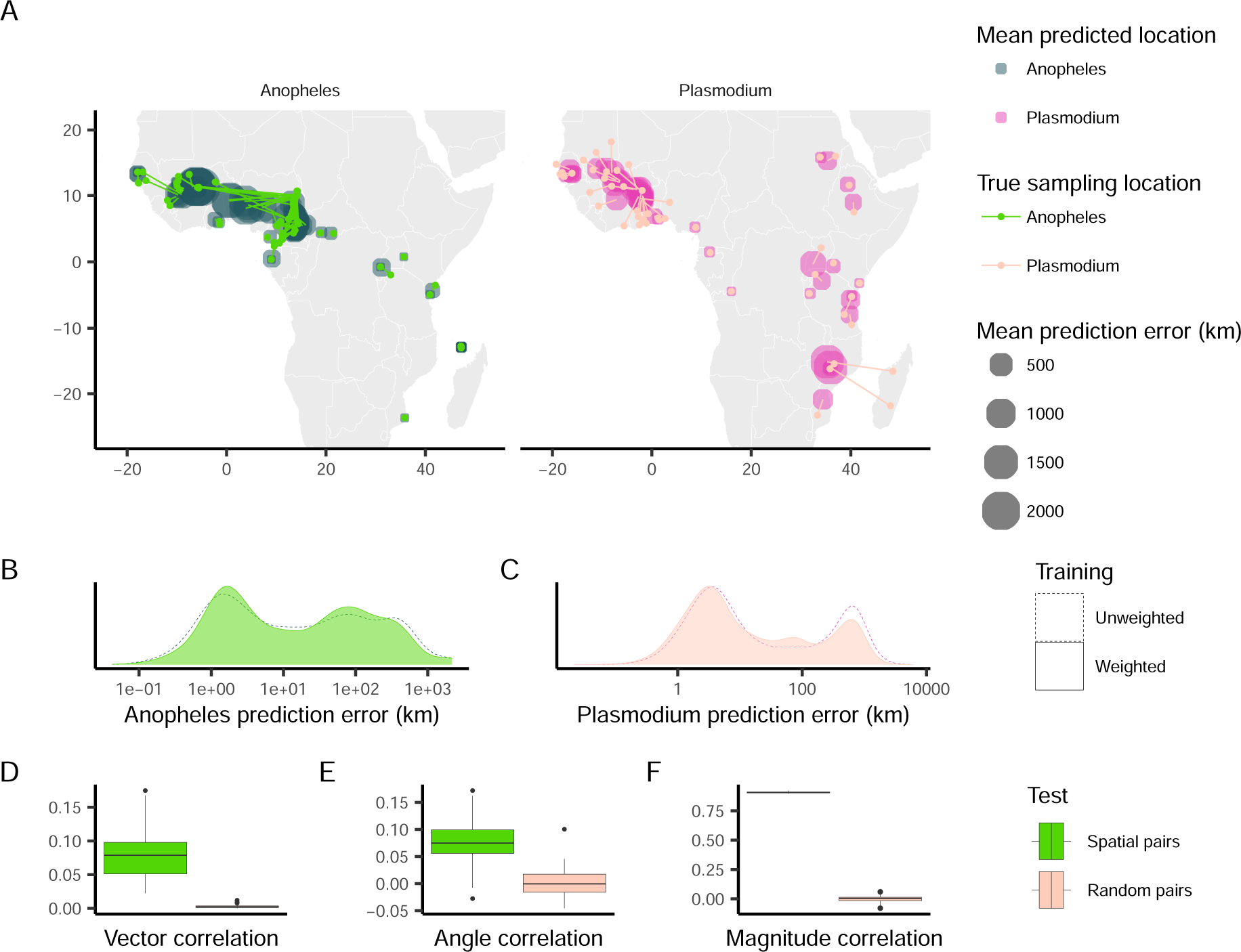
Predicted locations of *Anopheles gambiae* and *Plasmodium falciparum* samples after training using sample weights. (A) Geographic centroids of predictions for each unique sampling site. True sampling locations are connected to their mean predicted location; the size of predicted locations represents error in km. (B, C) Distribution of error magnitudes for both unweighted and weighted *A. gambiae* and *P. falciparum* predictions. (D, E, F) Vector, angle, and magnitude correlations between spatially paired or randomly sampled *A. gambiae* and *P. falciparum* weighted residuals.

Using sample weighting during training, we were able to reduce prediction error in our empirical applications, and likely in turn reduce the degree of spatial bias in predictions. Even so, the resulting prediction errors when applying weighted Locator to *A. gambiae* and *P. falciparum* populations still display similar patterns and are significantly correlated between the two species. This indicates that while the spatial distribution of sampling locations may contribute to prediction errors, Locator is nonetheless influenced by genetic signals of dispersal.

## Discussion

Motivated by the observation that the residuals in spatial predictions from population-scale genomic samples of *Anopheles gambiae* and *Plasmodium falciparum* are correlated, here we ask to what degree is this correlation the biological signal of the linked evolutionary history between host and parasite versus an artifact of spatially biased sampling? Battey et al. (2020a) showed that in some well-documented cases, large Locator residuals were associated with known admixture events or population movements. Because *Plasmodium* lives a portion of its life cycle in an *Anopheles* host, the geographic histories of these two species are correlated by necessity, thus spatial assignment predictions from genetic variation would be expected co-vary between host and parasite. While that is so, the spatial sampling of genotypes across the landscape is highly non-uniform, even with the massive publicly available datasets of the Ag1000G and Pf7k consortium projects. This of course makes sense – for instance some sampling locations will be more easily accessible than others – but to what degree will this impact our predictions? Here we use simulation to explore the impact of spatial sampling bias on the correlation of spatial assignment predictions between host and parasite. We find that the correlation of spatial assignment predictions between *A. gambiae* and *P. falciparum* is sensitive to the spatial sampling bias of the host, that biased spatial sampling leads to biased predictions, and demonstrate conditions under which we may or may not expect geographic bias in prediction error. We then examine how the effect of spatially biased training data might be mitigated and evaluate alternative strategies towards this end. Finally, by implementing algorithmic improvements to Locator and comparing spatially explicit simulations to empirical data, we show that the magnitude and degree of residual correlation between *A. gambiae* and *P. falciparum* significantly exceeds that expected from algorithmic bias, identifying an instance of co-geographic patterns of genetic variation between parasite and host.

### Sample weights reduce biased inference

A fundamental problem in statistical modeling is that biases in training data can easily lead to biased predictions. This is true of all methods to various degrees, but it could be argued that supervised learning approaches might be expected *a priori* to be more susceptible. Given the increasing ubiquity of supervised machine learning in society, as well as its growing popularity in genetics applications, it is important to understand the specific ways in which biased training data can affect predictions and how best to deal with them (e.g., Buolamwini and Gebru, 2018). Here, we examined Locator’s predictions from spatially biased training data and demonstrated methods that can reduce prediction bias caused by these training sets. Unsurprisingly, sample weighting during training was more effective at reducing bias than simply downsampling data; this makes sense as sample weighting still exposes the network to all available training samples. However our methods of sample weighting did not fully eliminate prediction bias, and in many regions of parameter space had either negligible effects on performance or made performance worse (Figure S5). While we tested a number of methods before settling on kernel density estimates, exploring more sophisticated applications such as using an adaptive bandwidth for the kernel density estimate or using a different function to scale sample weights may further improve accuracy and generalizability. Additionally, investigating the effects of sample weighting on different spatial sampling sampling schemes, such as point or midpoint sampling, would be informative as to how the method performs on empirical datasets.

We have described a method that can reduce prediction bias when training with unbalanced datasets, but this solution is ultimately limited by the bias-variance tradeoff. In our case, sample weighting successfully reduced both error magnitude and bias in our simulated datasets, however the overall variance in these values is greater than when training without sample weights. At its core, we are running up against a inferential limitation of our model. As we try to make our model more generalized - training Locator to predict locations across the entirety of the landscape - we sacrifice the precision of our inference. Ultimately, and like all inference methods, Locator will be limited by the distribution of data used during training. Spatial variation in training set density may represent the ecology of the organism: undersampled regions may reflect local changes in population density and regions with an absence of samples may represent locations from which the species is excluded. On the other hand sparsely-sampled portions of the landscape may occur due to simple sampling biases, such as unequal sampling effort, and thus pose a statistical challenge to correct for. Our implementation of sample weights has the power to increase prediction accuracy and reduce the bias that results from spatially imbalanced training data, but like in other biased inference problems, performance, parameterization, and interpretation of predictions are dependent on the given dataset.

### Empirical residuals reflect signal beyond sampling bias

In this study, we carefully assessed the potential for prediction bias in spatial genotype assessments of our empirical datasets. Using a novel modification to our methods and spatially explicit simulation, we demonstrated that the correlation between *A. gambiae* and *P. falciparum* prediction errors in Sub-Saharan Africa is significantly greater than we would expect in comparison to a simple null distribution This correlation in patterns of error persisted even after applying sample weights and improving overall prediction accuracy. Further, comparing empirical *A. gambiae* predictions to those from forward-in-time, continuous space population genetic simulations meant to mimic mosquito population dynamics, we demonstrated that the correlated residuals we observe in *A. gambiae* and *P. falciparum* predictions can not be explained by sampling alone. Although our model of *A. gambiae* is relatively unsophisticated, and does not incorporate major features of population ecology and evolution such as habitat variation, local adaptation or segregating chromosomal inversions, we have no reason to suspect that including these features would affect our results. In comparison to empirical *A. gambiae* predictions, residuals from our our simulated data were unimodal and generally smaller than the empirical residuals. This is particularly noticeable for individuals from West Africa, where the residuals are systematically larger than what we observe in the rest of the data set and suggest high migration rates and source-sink dynamics over that portion of the range in agreement with previous observation (e.g. Clarkson et al., 2020). While our simulated *A. gambiae* model provides a decent baseline for comparison, we are not directly modeling the linked life-histories of host and parasite. A more complex, two-species simulation, which models the interaction between *A. gambiae* and *P. falciparum* could help to reveal the degree to which host-parasite co-dispersal influences shared patterns of gene flow and how these patterns can be detected using inference methods like Locator.

We hypothesize that the correlation we observe here can be attributed to common patterns of dispersal between the two species - as *P. falciparum* depends on the *A. gambiae* mosquito for part of its life cycle, its movement will be partially limited by the movement of its mosquito host. The majority of this correlation structure seems to be driven by prediction errors in West Africa, where *A. gambiae* populations have been shown to have increased levels of gene flow and reduced differentiation (Clarkson et al., 2020). Indeed when applied to the Ag1000g dataset, Locator’s high prediction errors in West Africa and accurate predictions in Central and East Africa align with known patterns of spatial population structure in the *A. gambiae* species complex (Clarkson et al., 2020). Under the assumption of paired host-parasite dispersal, we would expect *P. falciparum* gene flow to also be elevated in West Africa and that prediction errors would have similar structure to *A. gambiae*, just as we observe. Further, humans are a key third player in the *A. gambiae*-*P. falciparum* interaction, as humans are hosts to both species, and accordingly *P. falciparum* dispersal has been shown to correlate with human mobility (Wesolowski et al., 2012). West Africa is a region of high human mobility (Wesolowski et al., 2014) - given that *A. gambiae* and *P. falciparum* are both human parasites (though with different degrees of linked dispersal), these patterns of gene flow may in part reflect anthropogenic factors.

### Hosts and parasites share evolutionary history

The degree to which a parasite depends on its host for dispersal shapes the strength of shared evolutionary history among taxa. Over longer timescales host-dependant dispersal can lead to co-diversification or even co-speciation between host and parasite (reviewed in Althoff et al. (2014)). At shorter timescales, hosts and parasites might share features of genetic variation that are correlated, for instance reductions in variation due to shared population bottlenecks. Accordingly empirical population genomic surveys of parasite species have repeatedly recovered the signature of shared evolutionary history among host and parasite. For instance Doyle et al. (2022) showed that estimates of historical demography for the human whip worm, *Trichuris trichiura*, shared signatures of the human out-of-Africa bottleneck event. Similarly Small et al. (2019) convincingly demonstrated that the biogeographic history of the filarial parasite *Wuchereria bancrofti* was shaped by a series of human dispersal events throughout the Indo-pacific. Counter examples of course exist. When host and parasite dispersal are less tightly linked, such as for the widely distributed intracellular parasite Hamiltosporidium and its host Daphnia species, co-biogeography and co-demography are not apparent (Angst et al., 2022). We speculate that the correlated residuals on geographic prediction between *A. gambiae* and *P. falciparum* is the result of their shared evolutionary interaction as host and parasite. Indeed Plasmodium is host-dependant with respect to dispersal, so our *a priori* expectation should be that of co-geography. The timescale of the processes which we are observing are very roughly on the order of the population size number of generations or fewer - using supervised machine learning methods, we are able to identify an instance of co-geography at the population scale.

## Materials and Methods

### Empirical data

We applied Locator to whole-genome sequencing data from the MalariaGEN Ag1000G Project and Pf7k Project (Anopheles gambiae 1000 Genomes Consortium, 2017; Pearson et al., 2019). From the Ag1000G dataset, we used 2,416 field-collected mosquitoes from the *Anopheles gambiae/coluzzii* species complex in sub-Saharan Africa. Samples used in the analysis belonged to *Anopheles gambiae* based on ancestry informative markers (Anopheles gambiae 1000 Genomes Consortium, 2017). From the Pf7k dataset, we used 3,315 *Plasmodium falciparum* field-collected blood samples collected in sub-Saharan Africa. For the *A. gambiae* dataset, we ran Locator in a windowed analysis across all chromosomes, using 2Mbp windows that captured roughly 100,000 SNPs each. A random 10% of samples were held out during training and used as a test set. This was repeated until all samples had been used in the test set, thus obtaining predicted locations for all 2,416 samples. For the *P. falciparum* dataset, we ran Locator using 500 kb windows, again capturing roughly 100,000 SNPs each, and held out a random 5% of samples during training. This same process was repeated when running Locator with sample weights.

### Spatial simulations

Simulations used for assessing Locator’s performance on spatially imbalanced datasets were run under the continuous space model described in Battey et al. (2020b) using SLiM v3 (Haller and Messer, 2019). Individuals were simulated across a 50 *×* 50 unit square landscape with an expected population density of 5 individuals per unit area and a mean parent-offspring dispersal distance that varied between 0.4 and 1.0 units. Simulated individuals were diploid, with two copies of a 10^8^ bp chromosome that recombined at a rate of 10^-8^ per bp per generation. Simulations were run for 50,000 generations, at which point nearly all tree sequences had coalesced. After simulations were completed, the resulting tree sequence was recapitated and neutral mutations were added at a rate of 10^-8^ per bp per generation. Finally, location metadata and genotype calls were saved for a given simulation. Genotypes were output as VCFs using the .write vcf() function in tskit, then converted to Zarr format using scikit-allel.

### Biased training sets

To create spatial bias in training data, we sampled training sets from the previously-described 50 *×* 50 unit continuous-space simulations with increasing probability that a sampled individual would be from the right half of the landscape (ie, *x >* 25.0). We refer to this as *training set bias*, which ranged from 0.5 (uniformly sampled) to 0.9 (a given sample having a 0.9 chance of being from the right half of the landscape). Individuals used for prediction (which we refer to as the *test set*) were sampled uniformly, enabling us to assess the trained model’s accuracy across the entire 50 *×* 50 landscape.

To assess how this bias in training data affects Locator’s predictions, we applied this method to simulations with per-generation dispersal rates (referred to as *σ*) between 0.4 and 1.0 units. We used biased sampling to sample 450 individuals as a training set, then trained Locator on 100,000 randomly chosen SNPs from these samples. The trained network was then used to predict the locations of 50 new uniformly-sampled individuals from the dataset. We defined a residual as the vector between a test set individual’s true location and their predicted location. To quantify bias in residuals across a given Locator run, we separately assessed the mean *x* and *y* component of residual vectors. Because the prediction set is uniformly sampled from the landscape, these values will be distributed around zero across multiple runs if predictions are unbiased.

For a set of simulations where *σ* = 1.0, we ran Locator in a full-genome windowed analysis, using 2Mbp windows containing roughly 8,000 SNPs each. Alongside this, we varied the number of individuals in the test set between 50 and 500, preserving a 9:1 train-to-test ratio in these analyses. Here, we defined a residual as the vector between a test set individual’s true location and the geographic centroid of their windowed predictions. These runs were used to assess the impact of training set bias on individual-level prediction uncertainty.

### Modifications to training data

#### Downsampling

To downsample densely-sampled regions of the landscape, we relied on the Generalized Random Tessellation Stratified (GRTS) method (Stevens. and Olsen, 2004), implemented using the geosample Python library. This method splits the landscape recursively into quadrants, then randomly samples from each quadrant to create a spatially balanced dataset. In order to assess how downsampling can affect Locator’s performance on datasets that are unevenly sampled across space, we applied GRTS sampling to our skewed training sets, resulting in a spatially even subset of a skewed training set’s samples being used during training. The same test sets of individuals were used for prediction.

#### Sample weighting

Sample weighting scales each sample’s contribution to the overall loss function when fitting Locator to training data. Weights were implemented using Tensorflow’s built-in sample weight argument to model.fit().

To calculate sample weights, we used scikit-learn (Pedregosa et al., 2011) to produce a Gaussian kernel density estimate of the density of spatial locations of training data across the landscape. Two free parameters are involved in assigning sample weights: bandwidth and *λ*. Bandwidth is used when fitting the kernel density estimate, and *λ* is used to exponentially scale the weights: if the estimated density at the location of the *i^th^* sample is *P_i_*, its weight *W_i_*is

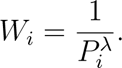

Bandwidth and *λ* values can be directly passed to Locator as a command-line argument or inferred using a grid search within Locator.

In our analysis, we used bandwidth values uniformly distributed between 0.25 and 5, and *λ* values ranging from 10*^−^*^5^ to 10.0. Methods to calculate sample weights using kernel density estimates or 2D histograms are now included in Locator using the--weight samples argument. Sample weights can also be calculated externally and provided as a TSV file. Finally, bandwidth and *λ* parameters for calculation of kernel density estimates can be provided as command-line arguments or identified using the GridSearchCV function from sklearn.model selection (Pedregosa et al., 2011).

### *A. gambiae* simulations

Populations used for assessing Locator’s performance on empirical *A. gambiae* data were also simulated using the same continuous space model in SLiM. Individuals were simulated across a map with boundaries representing the Sub-Saharan range of *A. gambiae* and that contained all empirical sampling locations. The landscape represented is 6900 kilometers across and 4750 kilometers high; to make this computationally feasible we reduced the scale of landscape and spatial interactions by a factor of 1/20.

An expected population density of 5 individuals per unit area was used in these simulations, resulting in a simulated population size of roughly 175,000 individuals. The effective population size of *Anopheles gambiae* is estimated to be 1,000,000 individuals, with a mutation rate of 3.5 *×* 10*^−^*^9^ and a recombination rate of 1.3 *×* 10*^−^*^8^ per base pair per generation (considering chromosome 2L) (Anopheles gambiae 1000 Genomes Consortium, 2017). These rates were scaled with the decreased population size in order to keep *θ* and *ρ* constant (*θ* = 4*Nµ, ρ* = 4*Nr*).

**Table 1:** Empirical estimates and scaled simulation parameters for *Anopheles* simulations.

|  | Empirical estimate | Scaled simulation |
| --- | --- | --- |
| Population size | 1000000 | 175000 |
| $\sigma$ | 13 km | 0.65 units |
| $\mu$ | $3.5 \times 10^{-9}$ | $2.5 \times 10^{-8}$ |
| $r$ | $1.3 \times 10^{-8}$ | $7.4 \times 10^{-8}$ |

We ran ten simulations and performed the same analysis on each. After running these simulations for 50,000 generations, we used coalescent simulation in msprime to recapitate the resulting tree sequences and added neutral mutations at a rate of 2.5 *×* 10*^−^*^8^ per bp per generation. To replicate the spatial distribution of empirical *A. gambiae* samples, we sampled an equivalent number of simulated individuals from each location where *A. gambiae* were sampled. Because multiple individuals in SLiM cannot occupy the exact same spatial coordinates, we used a k-nearest neighbors algorithm to sample simulated individuals around each empirical sampling point, then assigned those individuals’ locations to the empirical point they were sampled by. This method is similar to real-world sampling strategies, where individuals existing in a small geographic region are often assigned the same spatial coordinates during sample collection.

We applied Locator to these simulated datasets in the same fashion as our empirical analysis, holding out a random 10% of samples during training. We again ran a windowed analysis using 2 Mbp windows, which each captured roughly 80,000 SNPs across a 10^8^ bp genome, representing slightly less genomic information than the empirical *A. gambiae* data.

## Data availability

The code used to produce this analysis is available on GitHub: github.com/clararehmann/ locator-residuals. Locator is available at github.com/kern-lab/locator and now includes updated sample weighting methods.

## Acknowledgements

We thank CJ Battey, Scott Small, and Nate Pope for comments and suggestions during this work. This research was supported by NIH awards R35GM148253 and R01HG010774 to ADK.

## Supplementary material

### Windowed analysis and uncertainty

While our analysis focused on trends across individual predictions in a dataset, we were also interested in how estimated uncertainty is affected by training data. By visualizing the spatial spread of predictions using different portions of the genome, windowed analysis can be used to assess the uncertainty in individual-level predictions (Battey et al., 2020a). When examining the windowed predictions for individuals in our simulated datasets, we observed a negative correlation between local density of training data and the spatial spread of an individual’s predictions across genomic windows, with this trend becoming more pronounced as training sets became more imbalanced (Figure S4A). When trained on uniformly-sampled data (bias = 0.5) with a dispersal rate of 1.0, the relationship between spatial density of training data and spatial spread of windowed predictions for a test sample is weakly negatively correlated: *r* = *−.*10*, p* = 0.02*, n* = 500. With a training set bias of 0.9, the relationship is much stronger: *r* = *−.*67*, p* = 2.8 *×* 10^-65^*, n* = 500.

### Effect of training set size

We also assessed how increasing the number of training samples affects Locator’s performance on spatially imbalanced datasets (Figure S4, panels B-D). Initial analysis on randomly sampled training sets had indicated that 450 training samples were sufficient for accurate inference of location (Battey et al., 2020a), but this number of training samples did not provide the same results when sampled unevenly from the landscape. Unsurprisingly, increasing training set size reduced the effects of training set imbalance, even in populations undergoing a high rate of dispersal (1.0 units per generation). While residuals are still biased - correlation between x-axis error and training set skew remains significant with 4,500 training samples (*r* = 0.72*, p* = 3.92 *×* 10^-9^) - the magnitude of these errors is reduced, likely reflecting the overall increase in training data.

### Sample weighting hyperparameters

During our grid search, we observed an inverse relationship between the magnitude of band-width and *λ* parameters required to reduce residual bias when training with spatially imbalanced data. At high values of *λ*, low bandwidth resulted in reduced bias (as measured by error along the x-axis), while overall prediction accuracy remained unaffected (as measured by y-axis error and residual magnitude). As bandwidth was increased, the value of *λ* that resulted in reduced prediction bias while maintaining accuracy decreased consistently. These combinations form a diagonal across the parameter space, with Locator performing differently on either side of this threshold (Figure S5).

When both *λ* and bandwidth were low, sample weights were relatively uniform across the landscape, which had a negligible effect on training outcomes: in the space beneath optimal *λ*/bandwidth combinations, Locator’s predictions were similar to those from unweighted training runs (Figure S6 C). Above the optimal threshold, where both *λ* and bandwidth values are high, performance was more inconsistent: residuals were biased along the x-axis and y-axis, and overall error magnitude increased well beyond that observed when training without sample weights (Figure S6 E). With these parameter combinations, the majority of samples were assigned extremely low weights, causing Locator to only focus on a few samples near the landscape boundaries during training and predict samples to be in some landscape midpoint.

### Comparison to SPASIBA

Prior to our introduction of Locator, the state-of-the-art statistical method for geographic assignment using genotype data was perhaps SPASIBA (Guillot et al., 2015). SPASIBA uses a clever implementation of a geostatistical model which learns smooth surfaces of allele frequencies over a landscape, and then predicts locations of unlabeled samples by maximizing their likelihood on this surface. We were interested to compare the performance of SPASIBA and Locator when using spatially imbalanced training data, in particular because biased training sets may affect traditional statistical and machine learning methods to a greater or lesser extent.

In Figure S7 we show results from SPASIBA applied to training sets of 10,000 SNPs drawn from samples of 450 individuals. In some cases we were only able to execute seven SPASIBA analyses for a given training set bias, as compared to ten Locator analyses, because jobs were killed after 30 days of execution. As with Locator, SPASIBA’s predictions were biased towards the right half of the landscape when run on imbalanced training sets. However, SPASIBA’s predictions suffered significant collapse towards the center of the sampling density in some cases, and often covered a small fraction of the landscape, with test individuals across a run all predicted to be in only a few unique locations. These results suggest that SPASIBA may struggle to a greater extent than Locator in the face of imbalanced training data, and more generally that biased training sets can influence prediction in traditional statistical as well as machine learning settings.

---

**Figure S1:**
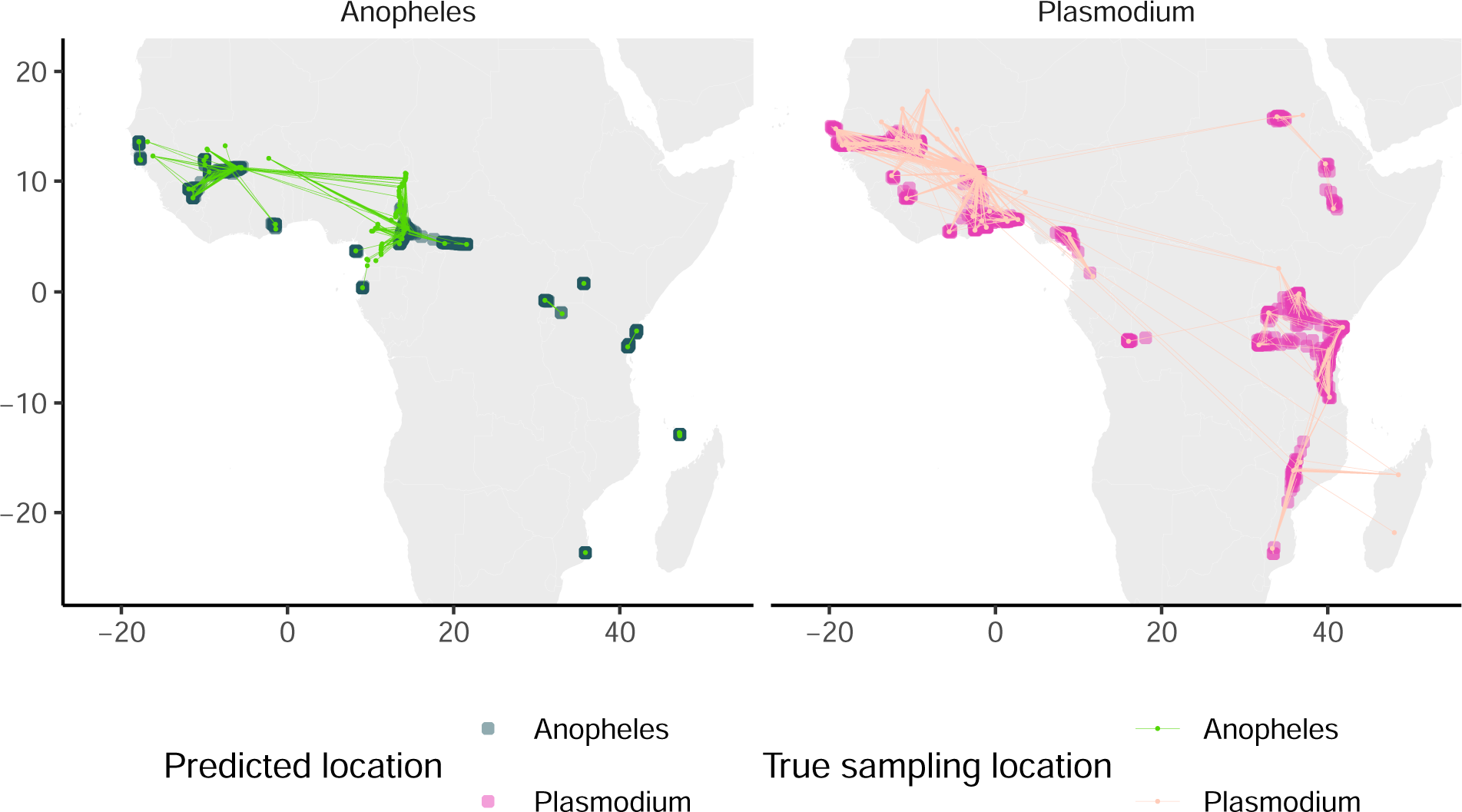
Predicted locations of all *A. gambiae* and *P. falciparum* samples. A sample’s true location is connected to the geographic centroid of its genome-wide windowed predictions.

**Figure S2:**
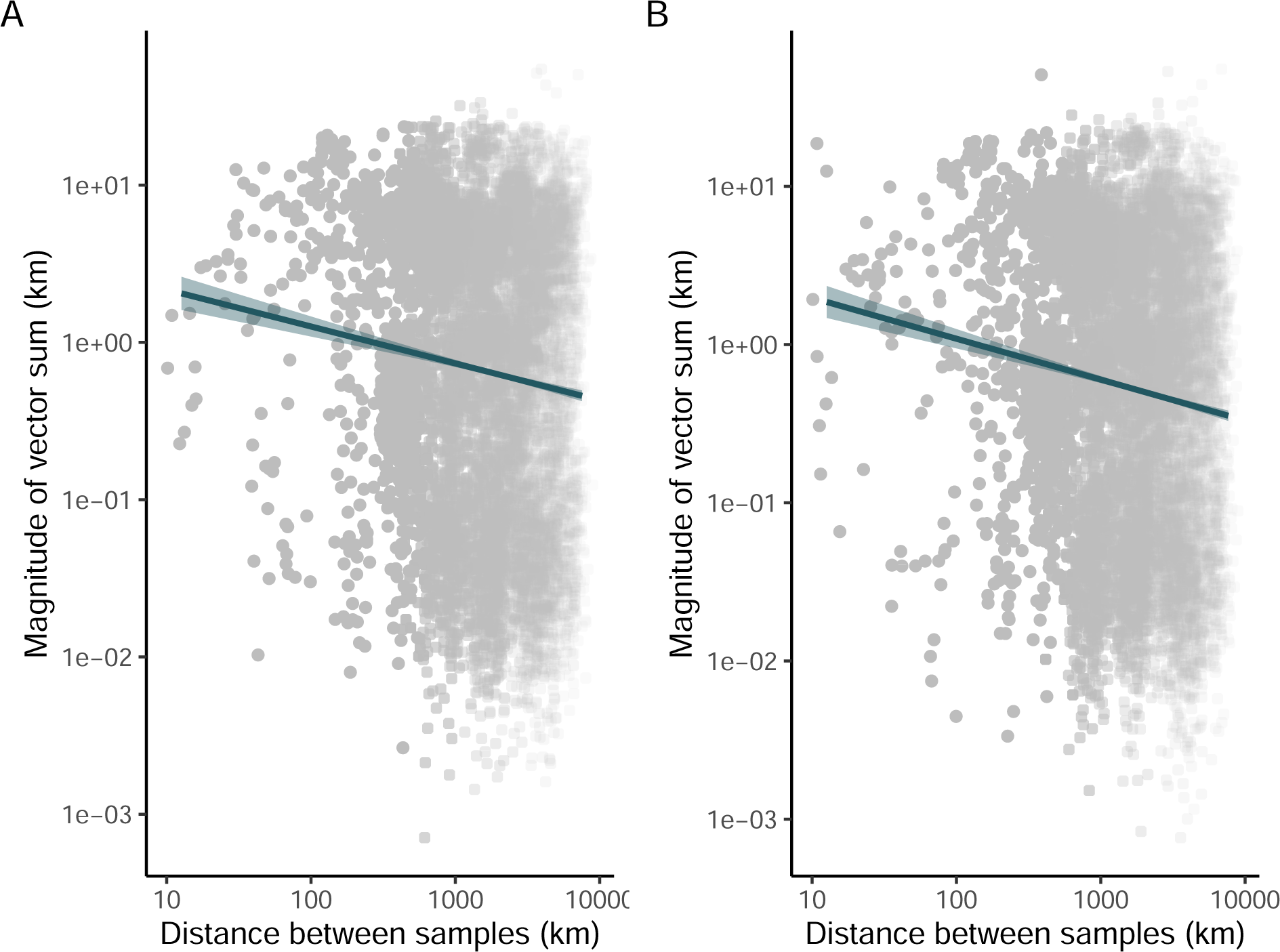
The magnitude of summed *A. gambiae* and *P. falciparum* residual vectors plotted against the distance between true sample locations. For both unweighted (A) and weighted (B) analyses, distance between samples is negatively correlated with the magnitude of summed residual vectors.

**Figure S3:**
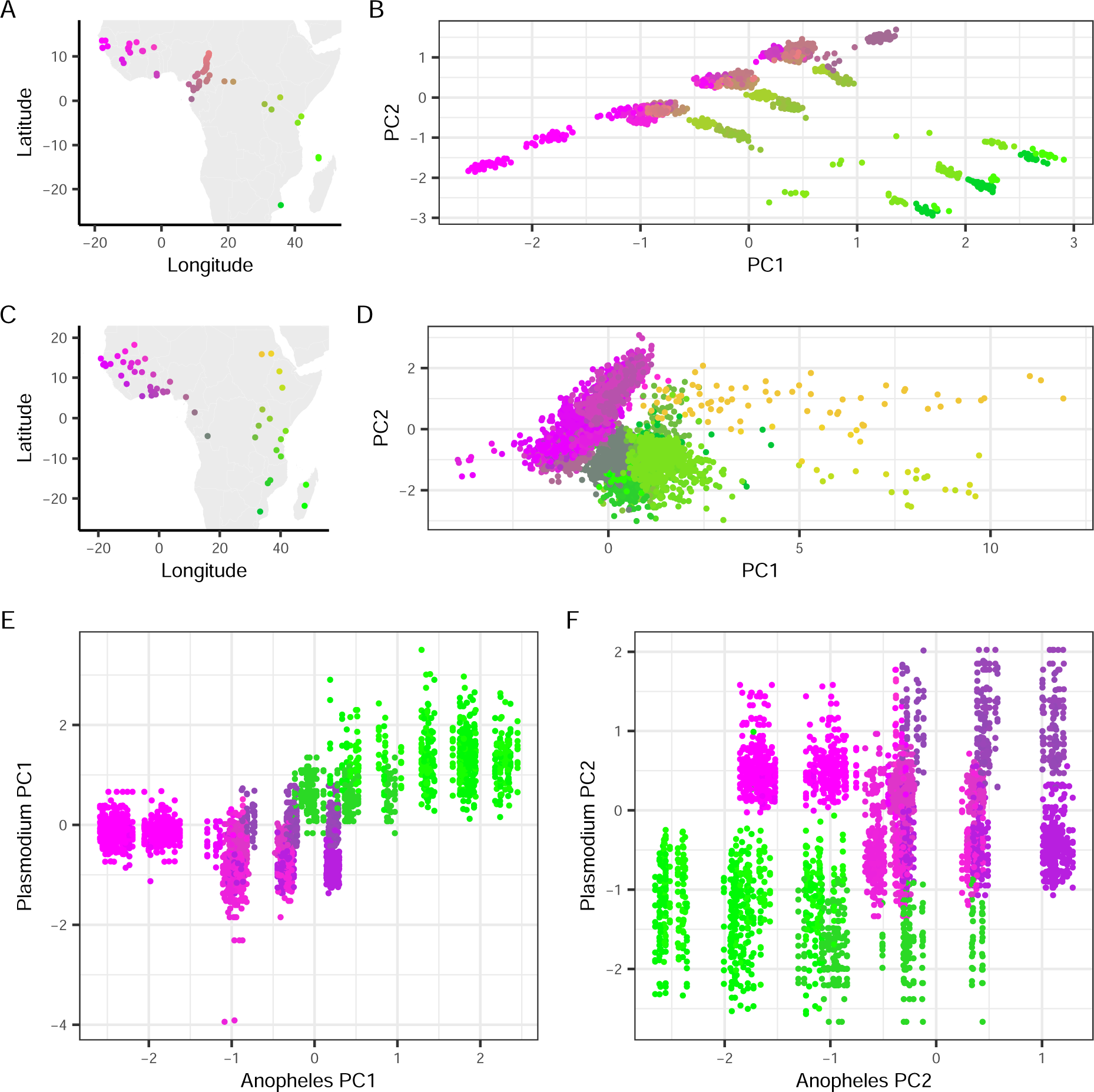
Spatially paired *A. gambiae* and *P. falciparum* principal components. (A) *A. gambiae* sampling locations and (B) principal components, with each sample colored by its spatial location. (C) *P. falciparum* sampling locations and (B) principal components, with each sample colored by its spatial location. (E) Relationship between the first principal component of spatially-paired *A. gambiae* and *P. falciparum* samples, with each pair colored by their spatial location. (F) Relationship between the second principal component of spatially-paired samples.

**Figure S4:**
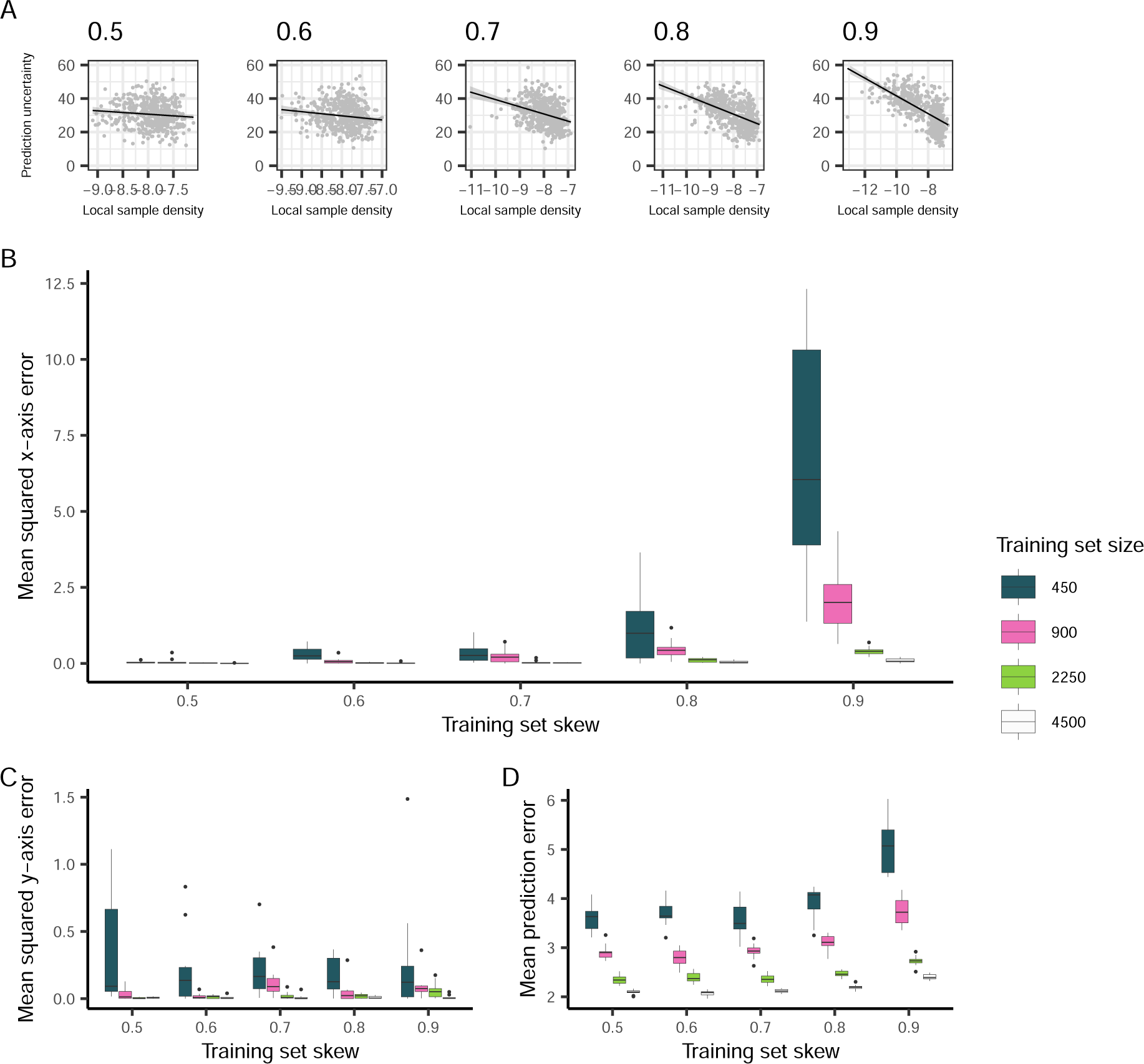
The effect of training set bias and size on Locator prediction error at a dispersal rate of *σ* = 1.0. (A) Spatial spread of per-individual windowed predictions as a function of local training set density at different levels of training set bias. (B) Mean per-run prediction error along the x axis as a function of training set bias, (C) mean per-run prediction error along the y axis, (D) mean per-run error.

**Figure S5:**
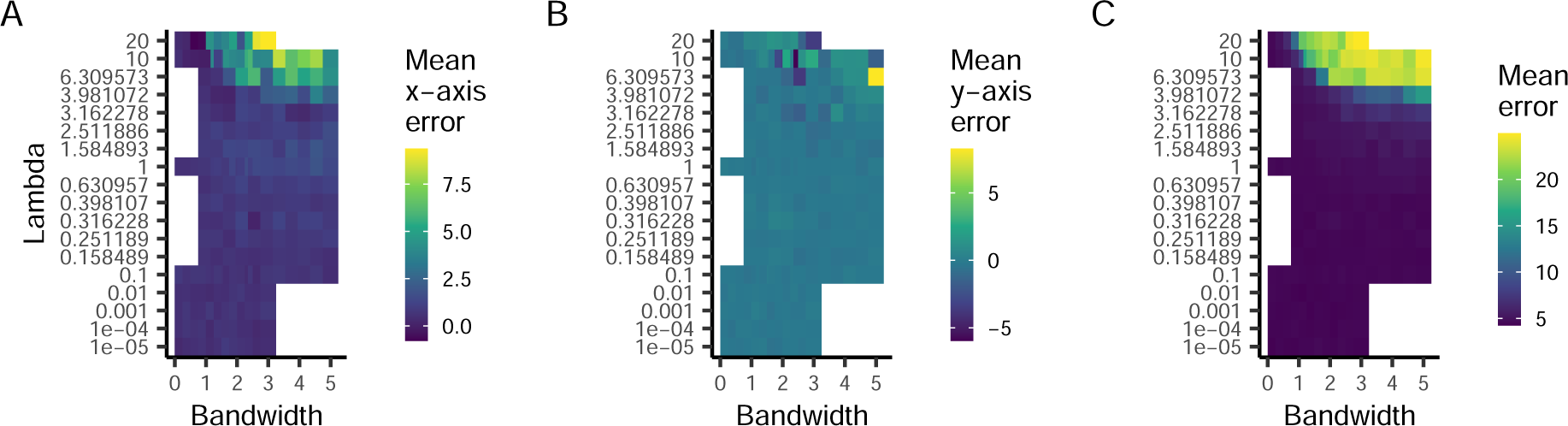
Results of grid search for *λ* and bandwidth parameters on *bias* = 0.9, *σ* = 1.0 training sets. (A) Mean x-axis error, (B) mean y-axis error, and (C) mean error for all tested combinations of*λ* and bandwidth.

**Figure S6:**
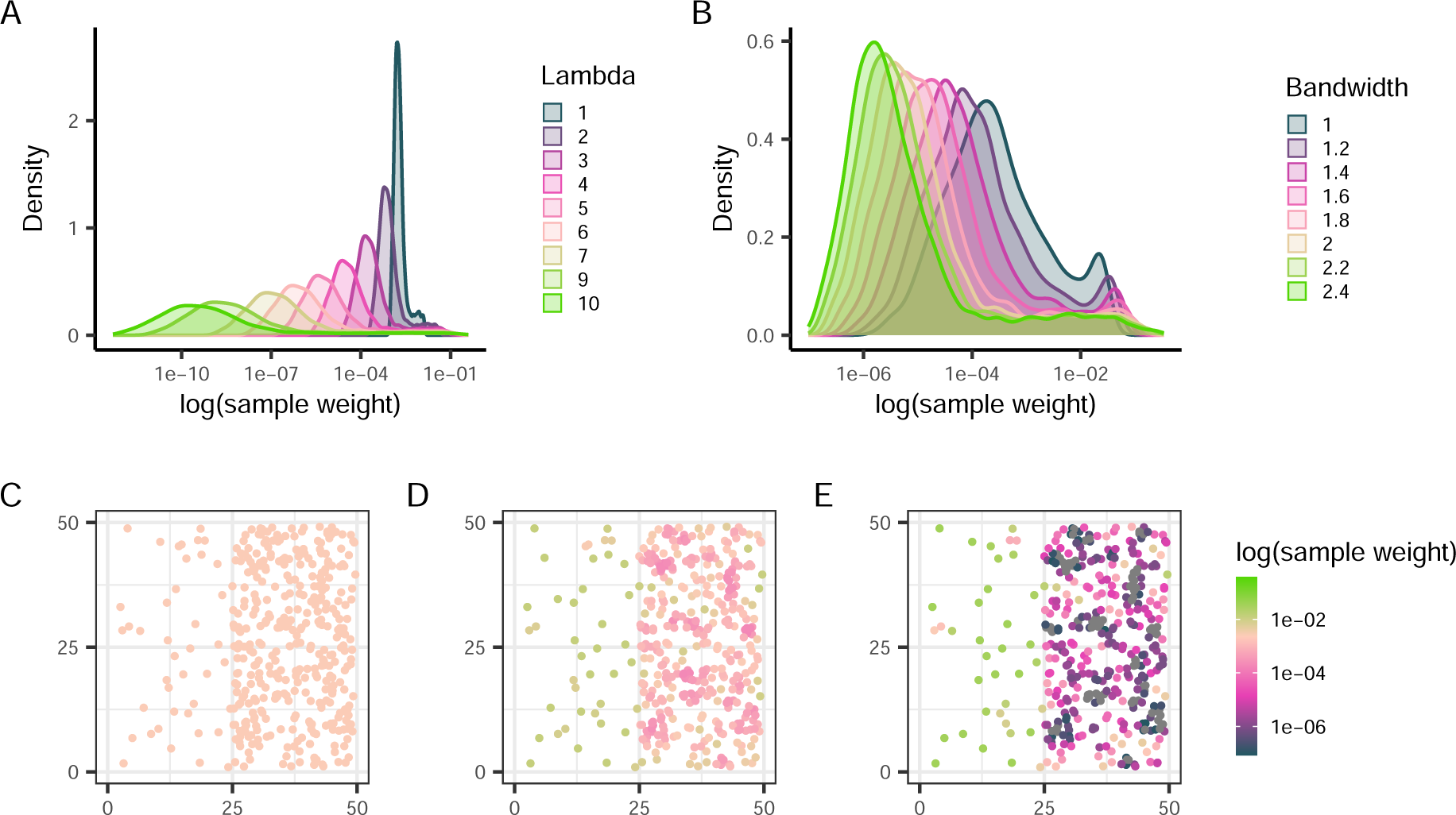
Bandwidth and *λ* parameters alter the distribution of sample weights. (A) Distribution of sample weights assigned to a given training set holding bandwidth constant and varying *λ*. (B) Distribution holding *λ* constant and varying bandwidth. (C, D, E) Biased training sets colored by their sample weights as *λ* increases.

**Figure S7:**
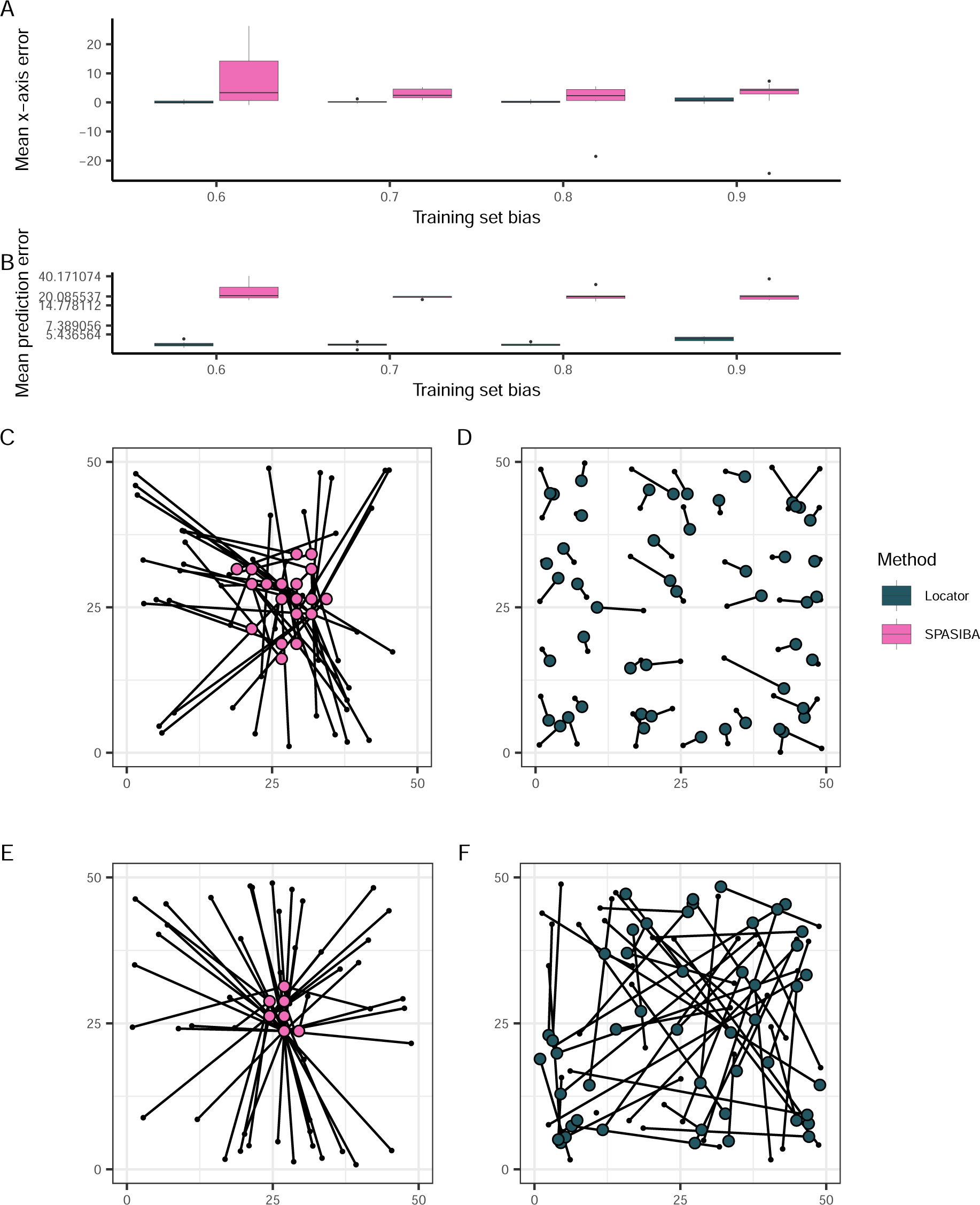
Predictions from biased training data using SPASIBA and Locator. Simulations had a per-generation dispersal rate of 1.0 units. (A) Mean x-axis error as a function of training set bias and (B) mean prediction error as a function of training set bias. (C, D) Example Locator and SPASIBA at *bias* = 0.5 and (E, F) *bias* = 0.9. True sample locations (black points) are connected to their predicted location (colored points).

